# Hmrbase2: A comprehensive database of hormones and their receptors

**DOI:** 10.1101/2023.01.31.526433

**Authors:** Dashleen Kaur, Akanksha Arora, Sumeet Patiyal, Gajendra Pal Singh Raghava

## Abstract

**Background and objective:** Hormones are essential for cell communication and hence regulate various physiological processes. The discrepancies in the hormones or their receptors can break this communication and cause major endocrinological disorders. It is, therefore, indispensable for the therapeutics and diagnostics of hormonal diseases.

**Methods:** We collected widespread information on peptide and non-peptide hormones and hormone receptors. The information was collected from HMDB, UniProt, HORDB, ENDONET, PubChem and literature.

**Results:** Hmrbase2 is an updated version of Hmrbase. The current version contains a total of 12056 entries which is more than twice the entries in the previous version. These include 7406, 753, and 3897 entries for peptide hormones, non-peptide hormones and hormone receptors, respectively, from 803 organisms compared to the 562 organisms in the previous version. The database also hosts 5662 hormone receptor pairs. The source organism, function, and subcellular location are provided for peptide hormones and receptors and properties like melting point; water solubility is provided for non-peptide hormones. Besides browsing and keyword search, an advanced search option has also been provided. Additionally, a similarity search module has been incorporated, enabling users to run similarity searches against peptide hormone sequences using BLAST and Smith-Waterman.

**Conclusions:** To make the database accessible to various users, we designed a user-friendly, responsive website that can be easily used on smartphones, tablets, and desktop computers. The updated database version, Hmrbase2, offers improved data content compared to the previous version. Homebase 2.0 is freely available at https://webs.iiitd.edu.in/raghava/hmrbase2.

## Introduction

Hormones are the key chemical players in coordinating activities between the cells and thus are important for maintaining and regulating homeostasis (1,2). These chemical messengers are secreted from specialised cells and undergo targeted signalling. The ability of the target cells to respond specifically to the hormones is possible due to receptors. These hormone-receptor complexes cause the derivation of intracellular consequences, which culminate in major molecular events leading to diverse cellular responses (3,4). These responses are essential in regulating physiological functions, including growth, development, metabolism and reproduction. The pictorial representation of hormonal response is shown in figure 1.

**Fig. 1.**
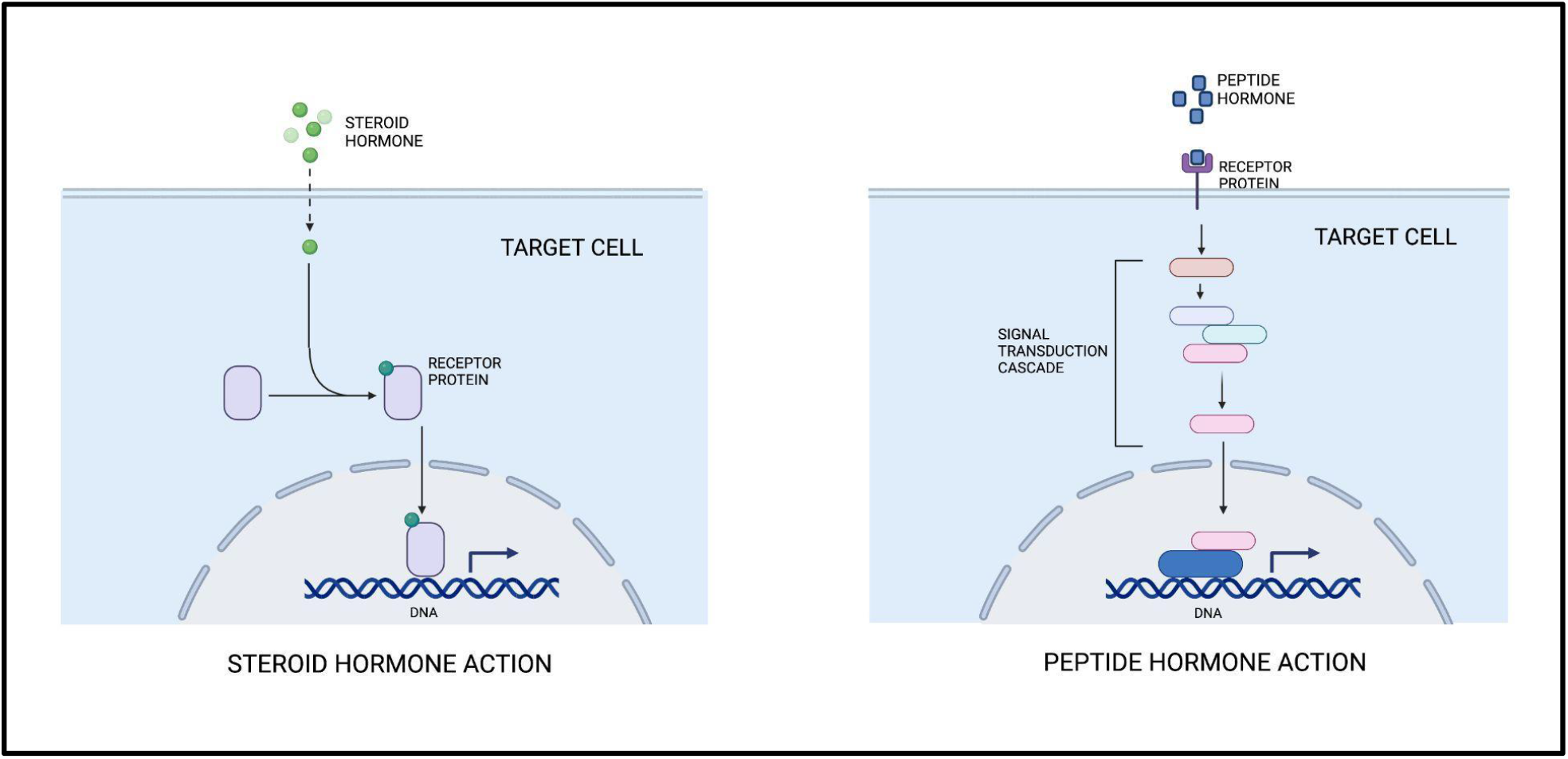
General representation of peptide and non-peptide hormone response

Hormones can be encoded by the genome (peptide-hormones), or they could be derived from other molecules (non-peptide hormones). Non-peptide hormones include steroid and amino acid molecules (5). They are usually produced in large quantities in plants and animals and transported to be received by the extracellular or intracellular receptors. A plethora of receptors is involved, including G-coupled, ion channels and nuclear receptors (6). Both plants and animals suffer from endocrine disorders due to excess and deficiencies of hormones. A comprehensive knowledge of hormones and their corresponding receptors will aid in diagnostics and the therapeutic interventions of endocrine disorders.

In recent years, there have been many databases related to receptors (like GPCRDB, BitterDB and NR-DBIND) (7–9) and a few related to ligands and their receptors (like PRRDB 2.0 and EndoNet) (10,11). PRRDB 2.0 is the database of immunological molecules. EndoNet is an information resource about intercellular regulatory communication. Thus, there hasn’t been a resource after the development of hmrbase which could provide comprehensive information on hormone and their receptors.

Hmrbase, a comprehensive database of hormones and their receptors, released its initial version in 2009. (12). Hmrbase contributed to the creation of new tools like SATPdb and FeptideDB (13,14). Since 2009, additional hormones and their receptors have been researched and identified, strengthening our understanding of endocrinology. The information regarding hormones and their receptors must therefore be updated in detail. Hmrbase2 is a revised and comprehensive re on hormones and their receptors. The new version contains in-depth details about hormones and receptors, including their localization and elaborate functions with information on the occurence, role, and their sequences. Additionally, the updated version contains information about the domains and pharmaceutical use of hormones which was missing in the first version.

The entries of hormones and their receptors are also linked to different databases like UniProt, PubChem, and PDB. This database also provides with sequence similarity search tool to help users map the query peptide sequences with hormonal peptide sequences. Thus, Hmrbase2 provides both comprehensive and easy-to-use information related to hormones and their receptors.

## Methods

### Data collection

The Hmrbase database, which was first created in 2009, has been updated to include new data on hormones (pharmaceutical use) and receptors that have been explored over the last thirteen years from many more organisms. This update reflects the growing interest in hormones across various species of organisms. There is a lot of information about peptide and non-peptide hormones and their receptors that can be found in various sources, including literature, databases, and online resources. To gather information about them, searches were conducted on web resources and databases using keywords related to hormones, including “hormone”, “phytohormone”, and hormone-receptors, including “hormone-receptors”, from the time period between May 2009 to November 2022. Information on peptide hormones and the hormone receptors was obtained from websites such as Uniprot (https://www.uniprot.org/), HorDB (http://hordb.cpu-bioinfor.org/), Endonet (http://endonet.sybig.de/) and Pubmed (https://pubmed.ncbi.nlm.nih.gov/) (11,15,16). To get information on non-peptide hormones, HMDB (https://hmdb.ca/) and PubChem (https://pubchem.ncbi.nlm.nih.gov/) were also used along with Endonet and Pubmed (17,18). Information was collected from various sources and was organised in tabulated form in the Hmrbase2 database. Redundant information was then eliminated.

### Database architecture and web interface

Linux-Apache-MySQL-PHP is the standard platform on which Hmrbase2 was created (LAMP). This database was created using Apache (version 2.4.7) as an HTTP server and MySQL (version 5.7.31). For producing responsive front ends that work with smartphones, tablets, and desktops, HTML (version 5), CSS (version 3), PHP (version 7.3.21), and Javascript (version 1.8) was used, and MySQL was used for building the back end. A standard interface was created using the PHP programming language. The working architecture of Hmrbase2 is explained in figure 2.

**Fig. 2.**
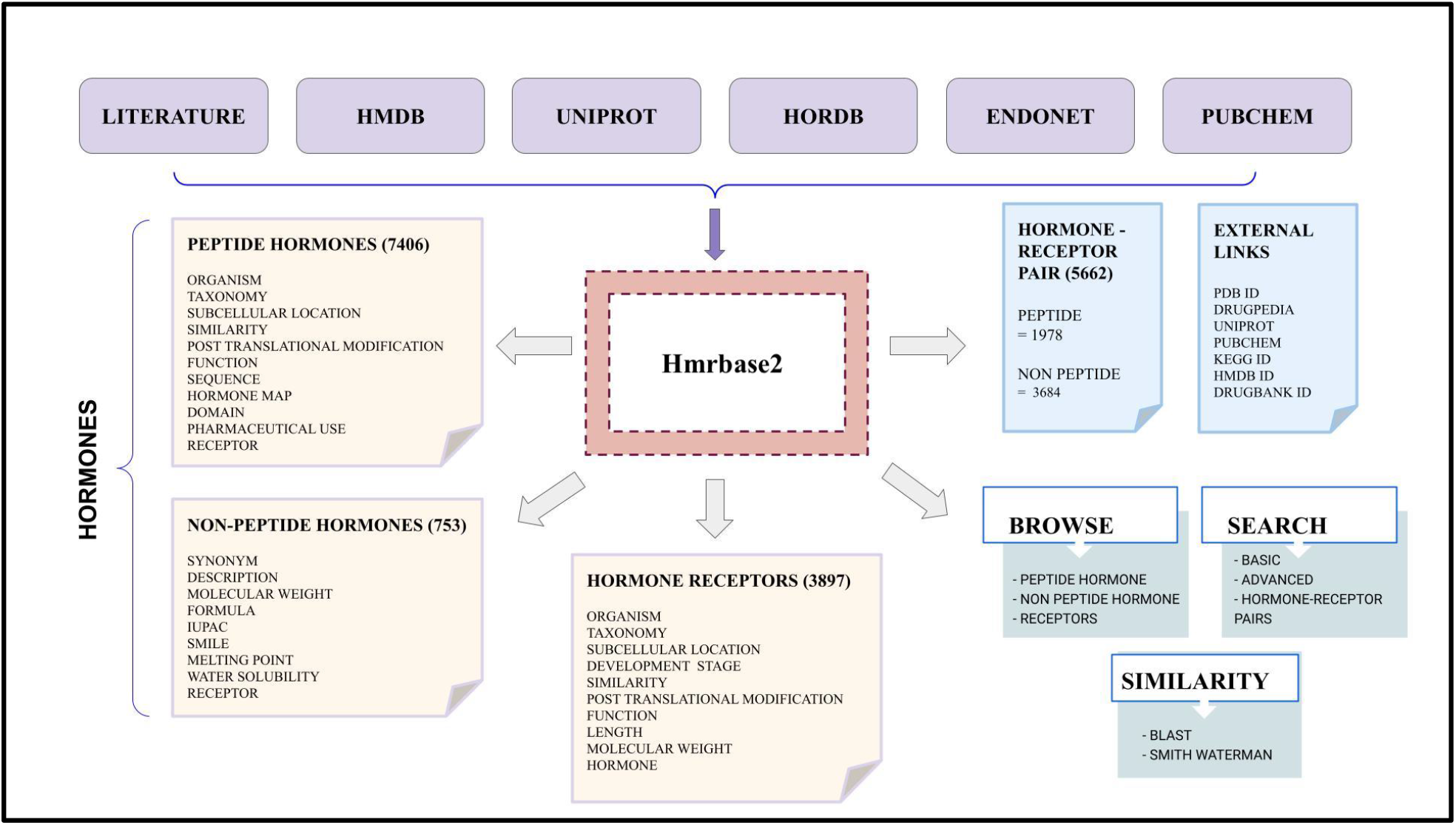
Architecture of Hmrbase2

### Data content

This database contains data about hormones and hormone receptors, including their sources, cellular, physical and functional characteristics, and sequences. The fields used to represent this information are particular to each type of hormone and receptor and include:

Peptide hormones containing the organism and taxonomy of the source organism, subcellular location, post-translational modification, function, sequence, receptor, and Uniprot ID; nonpeptide hormones having synonym, description, formula, IUPAC, SMILES, corresponding receptor, PDB ID, PubChem ID, KEGG ID, and HMDB ID; and receptors with organism and taxonomy of the source organism, subcellular location, post-translational modification, function, hormone, and Uniprot ID.

### Web interface

Hmrbase2 web server is openly available which is built to help the community. It provides widespread information about the hormones and their receptors on one platform. There are three significant modules included in the web server for an effortless search facility, i.e., ‘Browse,’ ‘Search,’ and ‘Similarity’.

#### Browse

The Hmrbase2 website has an uncomplicated and fast browsing option that makes it easier to get data from the database. Data can be explored based on fields specific for peptide hormones, non-peptide hormones and hormone receptors. This permits users to search through different categories of data even if they are unsure what they are looking for.

#### Search

There are two different search pages for basic and advanced searches. Besides that, the search page for hormone-receptor pairs has also been incorporated separately for peptide hormone-receptor pairs and non-peptide hormone-receptor pairs.

i. <u>Basic Search</u>: Separate Basic Search pages are provided for peptide hormones, non-peptide hormones and receptors. The architecture of all pages is identical except for the data field options. The searching here can be performed on an individual field or on all fields simultaneously, using a specific keyword to extract data from the database. The search option is available for all the data fields available in the database. Users can define which search results are to be displayed.
ii. <u>Advanced Search</u>: In advanced search, separate pages are provided for peptide hormones, non-peptide hormones and receptors. With additional boolean operators (such as AND, OR, and NOT), advanced search allows users to submit many queries at once and receive results accordingly. Users can download the displayed entries in comma-separated format.

#### Similarity

To enable similarity-based search, the Basic Local Alignment Search Tool (BLAST) and Smith-Waterman algorithms have been implemented (19,20). The user submits the FASTA format peptide sequence with default or specified settings to the BLAST module, and the server runs the BLAST search against the data stored in the database. Similarly, the Smith-Waterman algorithm searches for peptides based on similarity.

## Results

### Data analysis

The Hmrbase2, the latest version of the Hmrbase database, contains a total of 12,056 entries, including 7,406 entries for peptide hormones, 753 for non-peptide hormones, and 3,897 for hormone receptors. A comprehensive update on hormones and their receptors has been added. In addition to the 1955 hormones from the first edition of Hmrbase, we have included 6203 more hormones, bringing the total number of hormones to 8158. Similarly, Hmrbase2 includes 901 total hormone receptors researched in the last thirteen years, as well as 2996 receptors from the previous database, for a total of 3897 receptors.

The database contains hormones and receptors from 803 organisms across the animal and plant kingdom. Most of the hormones represent human hormones. The distribution of source organisms for both peptide hormones and receptors is depicted in figure 3 and proportion of peptide and non-peptide hormones present in this database is shown in figure 4. The amino acid sequences of mature peptide hormones and SMILES of non-peptide hormones are provided in the database. The entries in the database are linked with databases to get more information if needed. PDB IDs and Pubchem IDs are provided to get structural information. Uniprot ID is provided for the peptide hormones and the receptors to get information about them. Hmrbase2 also hosts 5662 hormone receptor pairs, 1978 for peptide hormones and 3684 for non-peptide hormones.

**Fig. 3.**
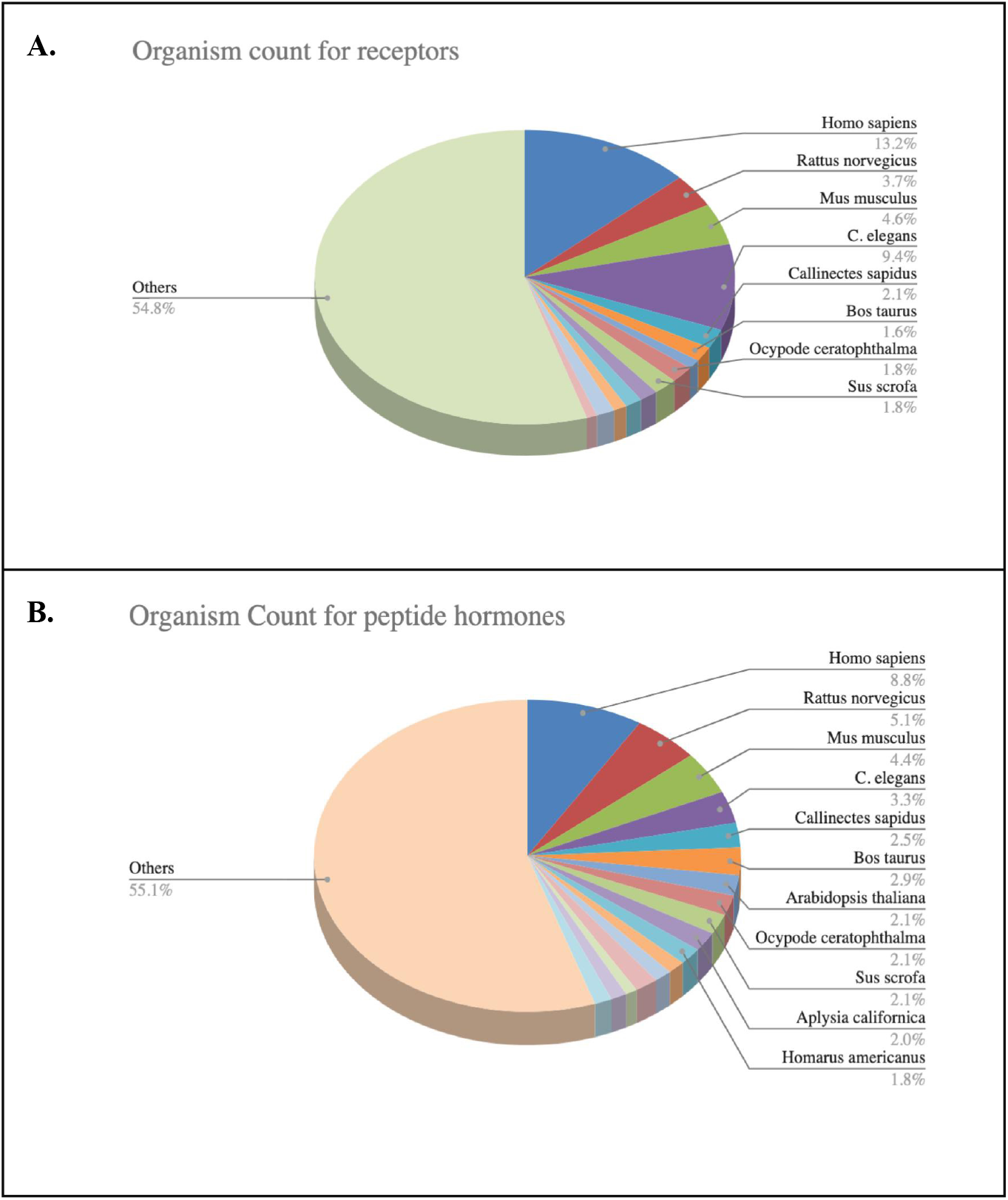
Distribution of source of peptide hormones (A) and hormone receptors (B).

**Fig. 4.**
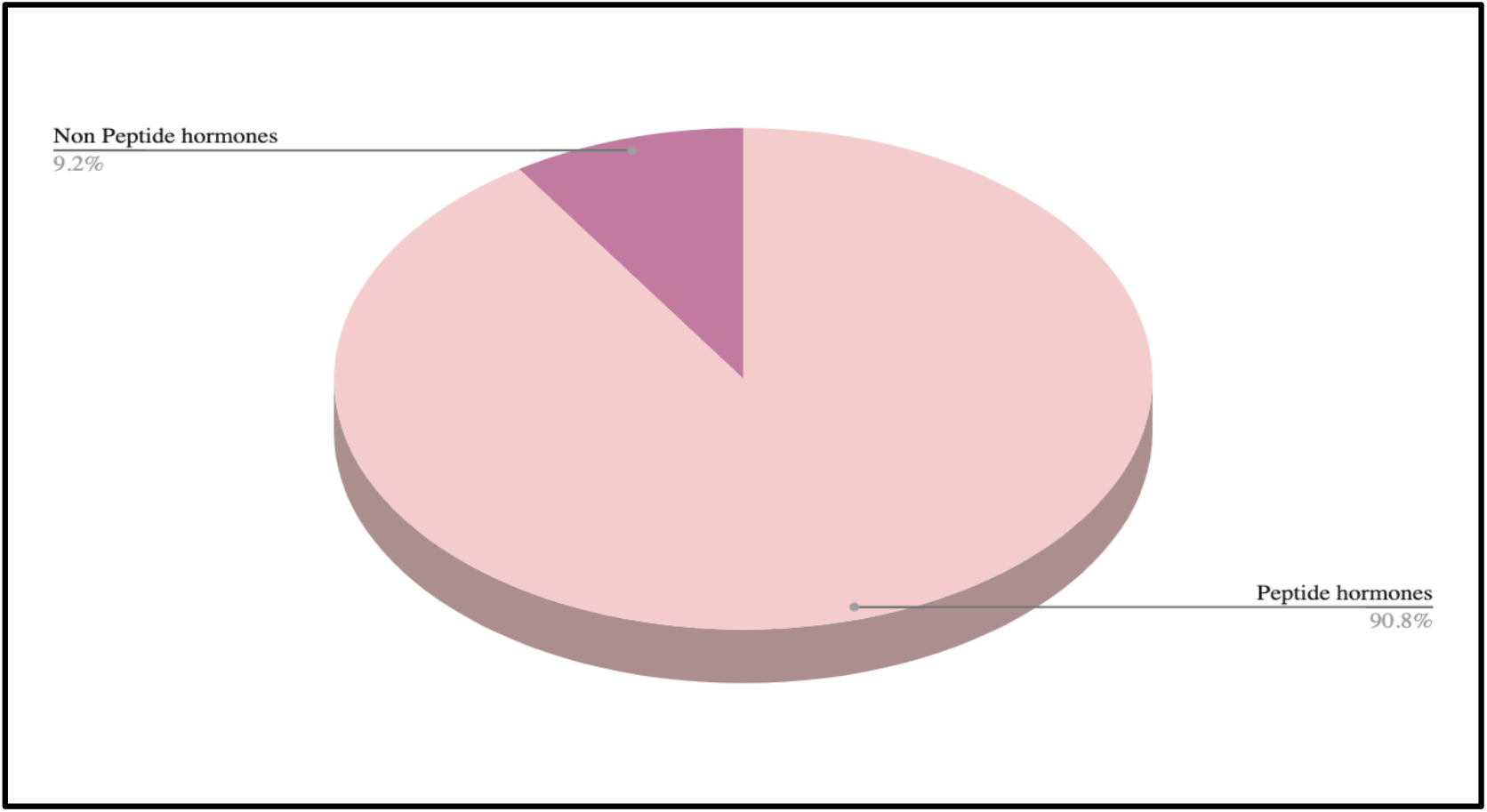
Proportion of peptide hormones and non-peptide hormones

### Comparison with the previous version

Hmrbase was developed in 2009 and comprised three different modules; one of the peptide hormones containing 1585 entries provides information such as their respective receptor’s name, sequence, sequence length, source organism and function. Another module is of non-peptide hormones containing 370 entries, providing information like name, function, IUPAC, SMILES and their receptor (12). There is also a module for receptors, containing information on 2996 receptors with their names, functions, length and post-translational modifications. In the updated version, we have included more hormones and their receptors, so the final data consist of 12056 entries. We have attempted to provide the pharmaceutical use of peptide hormones in addition to their name, type, source, and origin. The updated version also hosts 5662 hormone receptor pairs compared to the 4121 in the previous version. Comparative statistics are shown in Table 1. To add more, data has been linked to PubChem, UniProt, and PDB to provide users with for maximum information.

**Table 1.** Comparison of data in two versions of Hmrbase database

|  | Peptide hormone |  | Non-peptide hormone |  | Total |  |
| --- | --- | --- | --- | --- | --- | --- |
| Category | Hmrbase | Hmrbase2 | Hmrbase | Hmrbase2 | Hmrbase | Hmrbase2 |
| Hormone | 1585 | 7406 | 370 | 752 | 1955 | 8158 |
| Receptors | 828 | 1597 | 2168 | 2300 | 2996 | 3897 |
| Hormone-Receptor pairs | 569 | 1978 | 3552 | 3684 | 4121 | 5662 |

### Comparison with the other available databases

There are a limited number of databases which provide information about hormones and their receptors. The updated version of Hmrbase, Hmrbase2, is the only available database which gives comprehensive information about peptide hormones, non-peptide hormones and their receptors. HORDB is a database of peptide hormones maintained by Dr.Zheng’s team and contains only peptide hormones (16). Endonet focuses on the intracellular network of molecules (11). The complete comparison of Hmrbase2 with the existing resources is shown in Table 2. We also integrated the data of Endone into our database to provide users with information about all the hormones and their receptors. We have also added the peptide hormones from HorDB. The aim of Hmrbase2 is to make a comprehensive all-inclusive collection of all hormones and their receptors for the assistance of the scientific community working in this field.

**Table 2.** Comparison of Hmrbase2 with the existing resources

| Database | Peptide hormones | Non-peptide hormones | Hormone Receptors | Last updated |
| --- | --- | --- | --- | --- |
| **Hmrbase2** | **7406** | **752** | **3897** | **November 2022** |
| Hmrbase | 1585 | 370 | 2996 | 2009 |
| Endonet | 447 | 29 | 615 | 2014 |
| HorDB | 5729 | - | - | April 2022 |

**Fig. 5.**
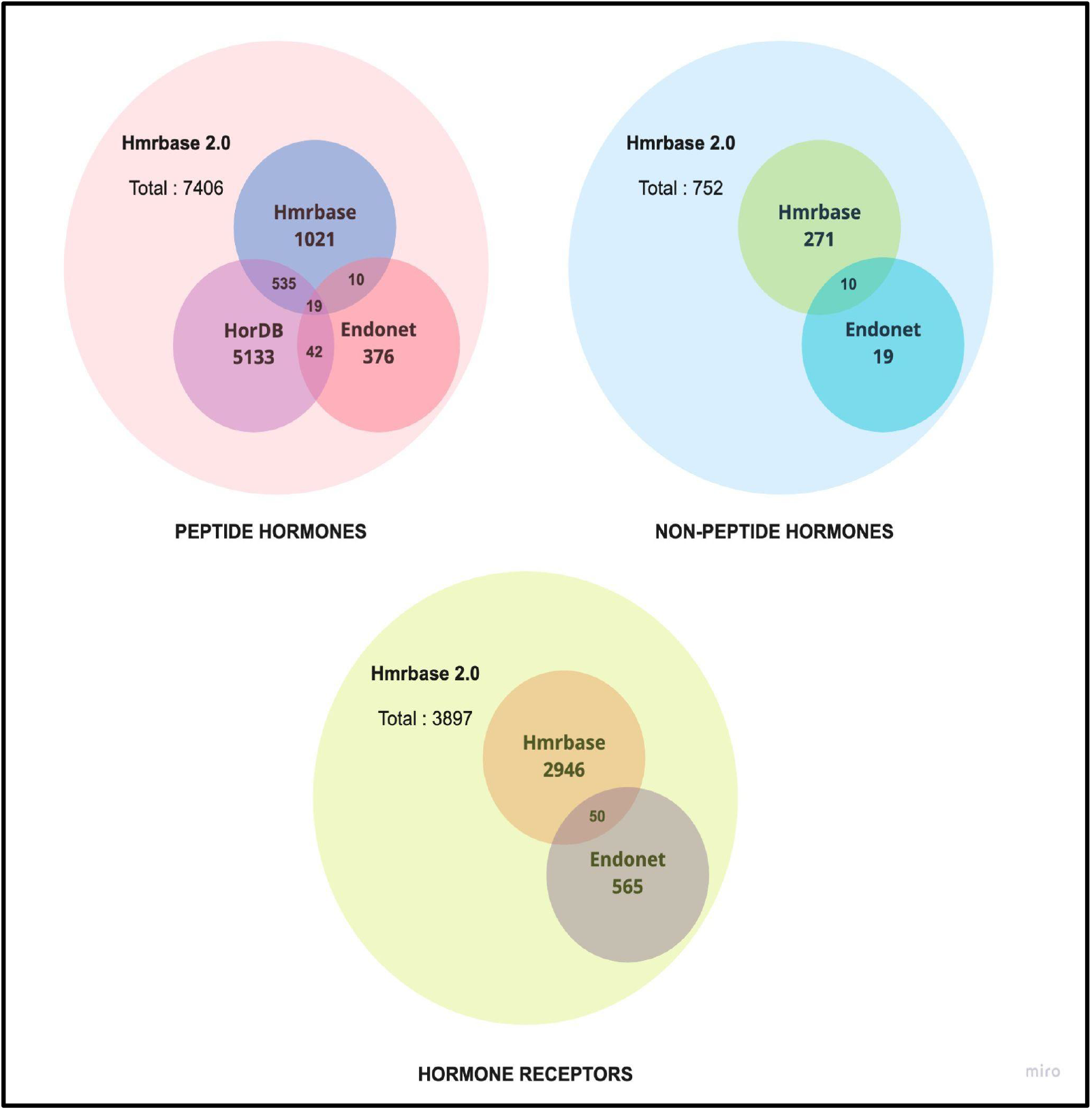
Information taken from various resources in Hmrbase2

## Discussion & Conclusion

Hmrbase2 provides a thorough knowledge source on hormones and their receptors. Bi-directional explanations of ligand-receptor interactions have been provided. Users have the option of beginning their search with the hormone entry (ies) and ending it with the appropriate receptor (s), and vice versa. The database will meet the needs of both theoretical and practising endocrinologists. The Hmrbase2 data structure is very user-friendly and straightforward. The mature hormone sequence is mapped onto its precursor protein sequence to characterise its functional modes.

Additionally, experimental scientists can use this knowledge to analyse the binding affinity of hormones to their respective receptors or to develop better ligands for a given receptor.

### The utility of database

Hmrbase2 can be used to get complete information about any hormone and its receptors on one platform. For eg., if a user is interested in Glucagon, a peptide hormone, one has to type Glucagon in the search bar given on the basic search page and select the required fields that user wishes to retrieve. The search results page and the other fields chosen will be displayed upon clicking the search button. Each entry is identified by a distinct ID that links to the hormone or receptor card for that entry, which provides detailed information about it.

### Limitation and update

Although we have made an effort to give comprehensive information on hormones and their receptors, some hormone and receptor sequences are not present in any chemical database or in the literature. Although the data is routinely filtered and carefully examined to reduce mistake, claiming complete accuracy would be unfair owing to potential human made errors. The first version of this database was published 13 years ago, but we will make an effort to revise the database regularly, ideally every three years.

## Author’s contributions

DK and AA collected the data. DK curated and analyzed the data. DK, SP, and AA developed backend of the webserver. DK and AA developed frontend of the webserver. DK, AA, and GPSR prepared the manuscript. GPSR conceived and coordinated the project. All authors have read and approved the final manuscript.

## Acknowledgements

Authors are thankful to Council of Scientific and Industrial Research (CSIR), Department of Biotechnology (DBT), Indraprastha Institute of Information Technology (IIITD) for fellowships and the financial support and Department of Computational Biology, IIITD, New Delhi for infrastructure and facilities. We thank the Department of Biotechnology (DBT) for providing an infrastructure grant to the institute.

## Data Availability Statement

All the datasets generated in this study are freely available at https://webs.iiitd.edu.in/raghava/hmrbase2/.

